# Brief optogenetic inhibition of rat lateral or ventrolateral periaqueductal gray augments the acquisition of Pavlovian fear conditioning

**DOI:** 10.1101/191841

**Authors:** Neda Assareh, Elena E. Bagley, Pascal Carrive, Gavan P. McNally

## Abstract

The midbrain periaqueductal gray (PAG) coordinates the expression and topography of defensive behaviors to threat and also plays an important role in Pavlovian fear learning itself. Whereas the role of PAG in expression of defensive behavior is well understood, the relationship between activity of PAG neurons and fear learning, the exact timing of PAG contributions to learning during the conditioning trial, and the contributions of different PAG columns to fear learning are poorly understood. We assessed the effects of optogenetic inhibition of lateral (LPAG) and ventrolateral (VLPAG) PAG neurons on fear learning. Using adenoassociated viral vectors expressing halorhodopsin (eNpHR3.0), we show that brief optogenetic inhibition of LPAG or VLPAG during delivery of the shock unconditioned stimulus (US) augments acquisition of contextual or cued fear conditioning and we also show that this inhibition augments post-encounter defensive responses to a non-noxious threat. Taken together, these results show that LPAG and VLPAG serve a key role in regulation of Pavlovian fear learning at the time of US delivery. These findings provide strong support for existing models which state that LPAG and VLPAG contribute to a fear prediction error signal determining variations in the effectiveness of the aversive US in supporting learning.

---

The midbrain periaqueductal gray (PAG) is divided into four columns located dorsomedial (DMPAG), dorsolateral (DLPAG), lateral (LPAG), and ventrolateral (VLPAG) to the cerebral aqueduct. It receives extensive descending projections from prefrontal cortex, extended amygdala, and ascending projections from spinal and trigeminal dorsal horn (Floyd, Price, Ferry, & Keay, 2006; Keay & Bandler, 2004; Rizvi, Ennis, Behbehani, & Shipley, 1991). The PAG, in turn, has ascending projections to hypothalamus, midline and intralaminar thalamus (Krout & Loewy, 2000), as well as descending projections to premotor and sensory regions in the brainstem and spinal cord (Keay & Bandler, 2004).

The PAG coordinates the expression and topography of defensive behaviors to threat (D. C. Blanchard, Williams, Lee, & Blanchard, 1981; Carrive, 1993; Depaulis & Bandler, 1991; Fanselow, 1991), with the DLPAG, DMPAG, LPAG associated with circa-strike defense, such as escape and flight, whereas LPAG and VLPAG are associated with post-encounter defense, such as freezing and quiescent immobility (Assareh, Sarrami, Carrive, & McNally, 2016; Bandler & Depaulis, 1991; Carrive, 1993; Fanselow, 1991). As such, the PAG is essential for expression of defensive behaviours as conditioned responses to a Pavlovian fear conditioned stimulus (CS) (Fanselow, 1991).

PAG also plays an important role in Pavlovian fear learning itself. Single unit recordings show variations in activity of DMPAG and DLPAG neurons to the aversive unconditioned stimulus (US) and CS during learning (Johansen, Tarpley, Ledoux, & Blair, 2010; Ozawa et al., 2016). Moreover, pharmacological manipulations of the PAG affect the acquisition, extinction, as well as second-order conditioning of fear (Cole & McNally, 2007; 2008; McNally & Cole, 2006; McNally & Westbrook, 2006; McNally, Johansen, & Blair, 2011). However, much remains to learned about this role for PAG in fear learning. It is known that PAG contributions to fear learning depend on the actions of endogenous opioids at the mu-opioid receptor in the LPAG and VLPAG (McNally & Cole, 2006; McNally, Pigg, & Weidemann, 2004b) and are also linked to activity in amygdala inputs, specially to DLPAG and DMPAG (Ozawa et al., 2016). However, the relationship between activity of PAG neurons and fear learning, the exact timing of PAG contributions to learning during the conditioning trial, and the contributions of different PAG columns to fear learning are poorly understood.

Here we assessed the roles of LPAG and VLPAG in Pavlovian fear conditioning. We used contextual and auditory cue fear conditioning designs to assess the effects of brief, optogenetic inhibition of LPAG and VLPAG neurons during the aversive US. The question of interest was whether optogenetic inhibition would affect fear learning. We also assessed the effects of this optogenetic inhibition on responding to a non-noxious visual threat - a looming stimulus – that provokes defensive responses in rodents (De Franceschi, Vivattanasarn, Saleem, & Solomon, 2016) as well as humans (Coker-Appiah et al., 2013) and causes strong recruitment of amygdala and PAG (Coker-Appiah et al., 2013; Wei et al., 2015; Yilmaz & Meister, 2013). The question of interest was whether optogenetic inhibition would affect responding to this non-noxious but threatening visual stimulus.

## Methods

### Subjects

Subjects were 135 experimentally naive adult male Sprague-Dawley rats (280-400g) housed in groups of four in a colony room maintained on a 12:12hr light-dark cycle (lights on at 7am). All procedures were conducted during the light phase. The procedures were approved by the Animal Care and Ethics Committee at the University of New South Wales.

### Apparatus

For contextual and cued fear conditioning, the chambers measured 24cm (length) x 30cm (width) x 21cm (height). The floors consisted of steel rods 4mm in diameter, 15mm apart. The top and rear walls, and front hinged door were made of clear Perspex. The end walls were stainless steel panels. The chambers were illuminated during testing with a house light located in the rear wall. They were situated in sound-attenuating cubicles (83cm length X 59cm width X 59cm height) with ventilation fans producing constant background noise. For auditory fear conditioning a 30s, 85dB [A scale] 2,800Hz tone was delivered through a speaker mounted to right side wall of the chamber. The US was a 0.7mA/0.5ms scrambled footshock.

For looming visual stimulus, a rectangular arena (50 x 50 x 50 cm [l x w x h]) was located in a sound attenuating chamber. A shelter (15 x 12 x 15 cm [l x w x h) was constructed along one wall. A 24” monitor positioned directly above the arena displayed a grey screen at all times other than during presentations of the looming stimulus. The looming stimulus was a black circle expanding from 1 cm to 25 cm diameter across 0.6 s controlled by a computer running MatLab^TM^ (Mathworks, Natick, MA).

For optogenetic inhibition, 8 - 9mW stimulation (measured as at the tip of an unimplanted fibre) was provided by 625nm LEDs (Doric Instruments, Quebec, Canada) controlled by a programmable driver (Doric Instruments, Quebec, Canada) and connected to the fibre optic cannula via 400μm/0.39NA patch cables.

### Surgery

Rats were treated with Carprofen (5 mg/kg), procaine penicillin (0.3 ml, 300 mg/ml), and cephazolin (0.3 ml, 100 mg/ml) prior to being anesthetized with ketamine/xylazine, and stereotaxically microinjected with 0.5 μl AAV5-hSyn-eYFP (3.7x10^12^ viral particles (vp)/ml titter) or AAV5-hSyn-eNpHR3.0-eYFP (2.7x10^12^ vp/ml) (UNC Vector Core) to target PAG neurons non-selectively. Vectors were infused unilaterally (3-min injection, 7-min diffusion) into LPAG (AP: -8.3, ML: -1.85, DV: -5.6) or VLPAG (AP: -8.3, ML: -1.85, DV: -6.6) followed by unilateral implantation of custom made fibre optic cannulas (400μm/0.39NA) (Sparta et al., 2011) targeting LPAG (AP:-8.3, ML: -1.85, DV: -5.3) or VLPAG (AP: -8.3, ML: -1.85, DV: -6.1) using standard stereotaxic technique. All coordinates are in mm bregma at 10° angle (Paxinos & Watson, 2007). Rats were allowed 21 days recovery prior to the start of the experiment.

### Procedure

*Contextual fear conditioning.* There were 4 groups: group eYFP (control) that expressed eYFP in LPAG or VLPAG; group LPAG that expressed eNpHR3.0 in LPAG; group VLPAG that expressed eNpHR3.0 in VLPAG; and group Offset that expressed eNpHR3.0 in LPAG or VLPAG. On Day 1, rats were exposed to the chamber for 5 min and tethered via patch cables. On Day 2, each rat received two 0.5s unsignalled footshocks 2 and 4 min after placement in the chamber and removed 1 min after the second footshock. eYFP, LPAG and VLAG groups received continuous photoinhibition during the 0.5s US whereas the offset group received these stimulations offset from the US (one 30 s after placement and one 3 min after placement). On Days 3 and 4, rats were returned to the apparatus and fear responses assessed for 10 min.

*Cued fear conditioning.* There were 2 groups: group eYFP that expressed eYFP in LPAG or VLPAG; group eNpHR3.0 (groups Halo) in LPAG or VLPAG. On Day 1, rats were placed in the conditioning chambers for 5 min while tethered via patch cable. Day 2, rats received 3 presentations of the auditory CS which co-terminated with the footshock US. Photoinhibition occurred during the 0.5 s US. The first trial commenced 2 min after placement in the chamber and remaining trials occurred at 2 min ITI. On Day 3 rats were tested for fear responses to the CS. They received 6 CS presentations commencing 5 min after placement in the chamber and with a 2 min intertrial interval (ITI).

*Looming threat*. There were 3 groups: group Loom - eYFP that expressed eYFP in LPAG or VLPAG; group Loom - Halo that expressed eNpHr3.0 in LPAG or VLPAG; and group Control – Halo that expressed eNpHr3.0 in LPAG or VLPAG. Food was removed from the animal cages 18hr before commencement of each day of experiment but was otherwise freely available. Rats received 4 days 20 min/day habituation to the arena and 15 Fruit Loops^TM^ were distributed randomly in the arena during every session to encourage exploration. Animals were habituated to the patch cable tethering procedure during the final two pre-exposure sessions. On test, rats were placed in the arena. 2 min later, the looming stimulus was presented. Photoinhibition overlapped (1 s prior, 0.6 s during, and 1 s after) presentations of the looming stimulus. There were three looming trials. The ITI varied with a minimum of 5 min since the last occurrence of any freezing and the rat must have been located in the centre of the arena for the next trial to begin. In the halo control group, the procedure (including photoinhibition) was the same but no looming stimulus was presented.

### Histology

After completion of the experiment, rats were deeply anesthetized with sodium pentobarbital (100 mg/kg i.p.) and perfused transcardially with 200 ml of 0.9% saline, containing 1% sodium nitrite and heparin (5,000 i.u./ml), followed by 400 ml of 4% paraformaldehyde in 0.1 M PB, pH 7.4. Brains were kept in the same fixative solution for 1 hr and then transferred to 20% sucrose solution overnight. Brains were processed for eYFP immunohistochemistry as described previously (Assareh et al., 2016). Sections were examined using an Olympus BX-53 Fluorescence microscope and placements determined according to the atlas of Paxinos and Watson (Paxinos & Watson, 2007). In order to be included in analyses, both eYFP and optic fibre placement had to be located in LPAG or VLPAG.

### Data Analyses

The amount of time spent freezing during context and/or CS presentations was scored every 2 s and converted to a percentage of observations. For looming, velocity pre, during and after presentations of the stimulus was assessed via Ethovision XT (Noldus, Wageningen, Netherlands) and the amount of time spent freezing in the 2 min after stimulus presentations was scored every 2 s and converted to a percentage of observations. These data were analysed via ANOVA testing orthogonal contrasts and the per contrast type 1 error rate (*α*) was controlled at the 0.05 level (Harris, 2004)

## Results

Figure 1A shows the extent of eYFP expression and location of the fibre optic cannulae for rats included in the analyses from contextual fear conditioning. Only animals with fibres and eYFP expressed in the appropriate PAG column were included. Inevitably there was spread of eNpHR3.0 and eYFP expression into the adjacent PAG column. Consistent with our past work, the location of the fibre was used to determine group assignment because we have shown that optogenetic manipulation of LPAG and VLPAG can have different effects on behaviour (Assareh *et al*., 2016). After histology, group sizes were: group eYFP n = 9, group Offset n = 9, group LPAG n = 7, and group VLPAG n = 7.

**Figure 1.**
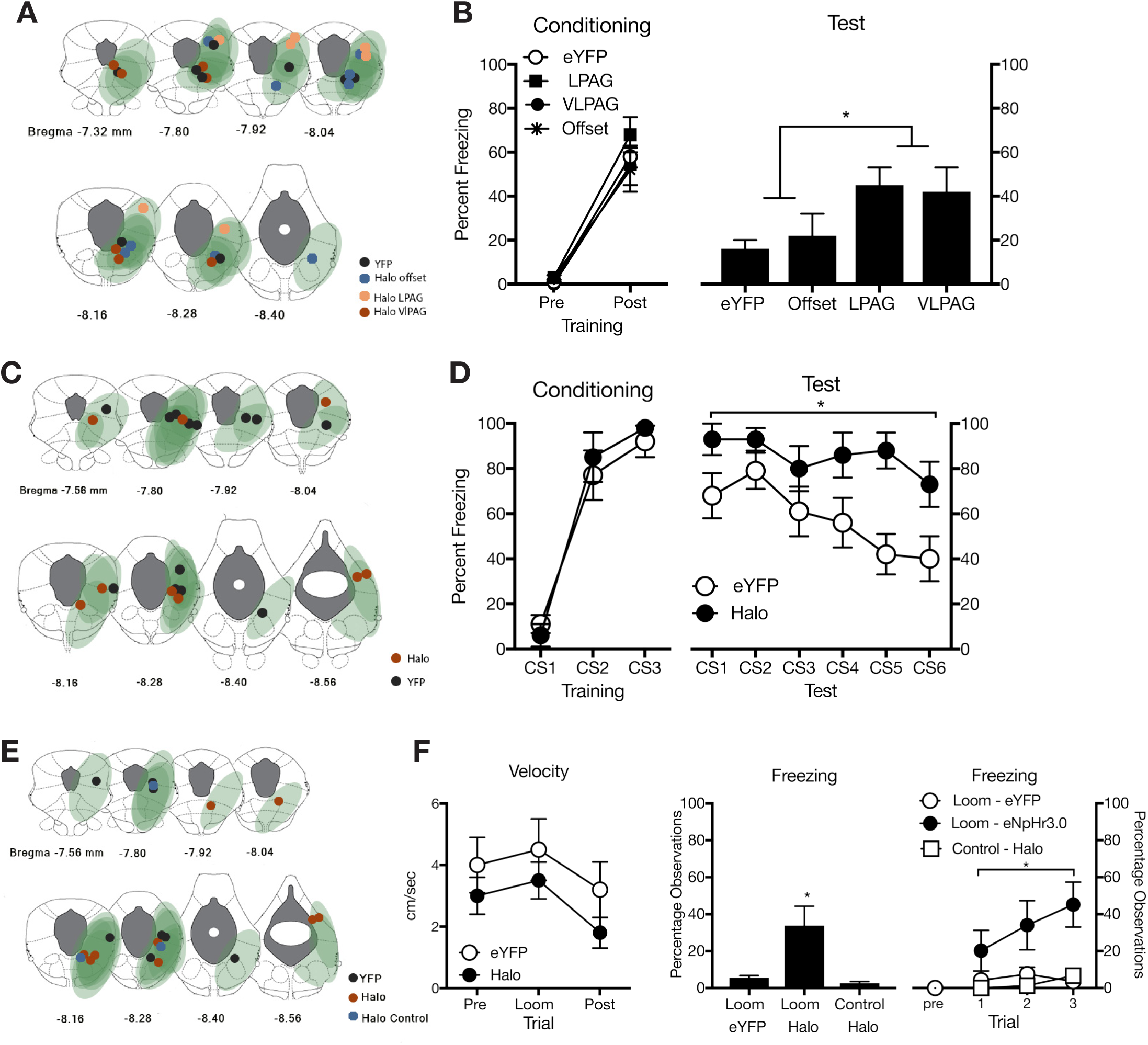
*A*, Extent of eYFP expression in PAG for each rat in the contextual fear conditioning experiment, with each rat represented at 10% opacity and location of fibre optic tips in LPAG or VLPAG. *B*, Mean (± SEM) levels of freezing during contextual fear conditioning and test (averaged across two test days). *C*, Extent of eYFP expression in PAG for each rat in the cued fear conditioning experiment, with each rat represented at 10% opacity and location of fibre optic tips in LPAG or VLPAG. *D*, Mean (± SEM) levels of freezing during cued fear conditioning and test. *E*, Extent of eYFP expression in PAG for each rat in the visual threat experiment, with each rat represented at 10% opacity and location of fibre optic tips in LPAG or VLPAG. *F*, Mean (± SEM) velocity during looming threat and mean (± SEM) levels of freezing in the 2 min post-looming stimulus. * p <.05.

The mean and standard error of the mean (SEM) levels of freezing during conditioning are shown in Figure 1B. During conditioning, there was significantly more freezing in the 3 min after the first shock than during the 2 min pre-shock period (F (1, 28) = 168.28, p < .0001). However, there was no overall difference in freezing between the LPAG and VLPAG groups versus the eYFP and Offset groups (F (1, 28) < 1, p > .05), between the LPAG versus VLPAG group (F (1, 28) < 1, p > .05) or between the eYFP versus Offset group (F (1, 28) < 1, p > .05). Moreover, there were no interactions between group and pre- versus post-shock periods (Fs (1, 28) < 1, p > .05).

The mean and SEM levels of freezing, averaged across the two test days, are shown in Figure 1B. There was significantly more freezing in the LPAG and VLPAG groups versus the eYFP and Offset groups (F (1, 28) = 6.71, p = 0.15) .05), but not between the LPAG versus VLPAG group (F (1, 28) < 1, p > .05) or between the eYFP versus Offset group (F (1, LPAG 28) < 1, p > .05). So, optogenetic inhibition of LPAG or VLPAG neurons during shock US delivery augmented contextual fear conditioning. This conclusion is unaltered if data from the first test (LPAG and VLPAG groups versus the eYFP and Offset groups, F (1, 28) = 6.47, p = 0.17; no other differences significant) or the second test (LPAG and VLPAG groups versus the eYFP and Offset groups, F (1, 28) = 5.61, p = 0.03; no other differences significant) are considered separately.

Figure 1C shows the extent of eYFP expression and location of the fibre optic cannulae for rats included in the analyses from cued conditioning. After histology, group sizes were: group eYFP n = 13, group Halo n = 9 (4 in LPAG and 5 in VLPAG).

The mean and SEM levels of freezing during conditioning are shown in Figure 1D. There were no differences in levels of pre-CS freezing (group eYFP mean = 5, SEM = 4; group Halo mean = 9, SEM = 4) (F (1, 20) < 1, p > .05). There was significant linear increase in CS-elicited freezing across conditioning trials (F (1, 20) = 232.10, p < .0001). There was no overall difference between groups in CS-elicited freezing during conditioning (F (1, 20) < 1, p > .05) and there was no interaction across trials (F (1, 20) < 1, p > .05).

The mean and SEM levels of freezing during test are shown in Figure 1D. For analyses we collapsed across LPAG and VLPAG halo groups because there was no difference between them in this or the preceding experiment. There were no differences in levels of pre-CS freezing (group eYFP mean = 45, SEM = 9; group Halo mean = 77, SEM =15) (F (1, 20) = 3.79, p > .05). There was significantly more freezing during CS presentations than during the pre-CS period (F (1, 20) = 7.82, p < .05). Overall, the Halo group showed significantly more CS-elicited freezing than the eYFP group (F (1, 20) = 5.60, p = .029). Freezing decrease significantly across the test trials (F (1, 20) = 17.91, p < .001), but this decrease did not differ between groups (F (1, 20) < 1, p > .05). The high levels of pre-CS freezing are problematic because they suggest greater contextual learning or generalization among the halorhodopsin groups. This is an important possibility, but it remains consistent with the finding that optogenetic inhibition of PAG neurons augmenting fear learning.

Figure 1E shows the extent of eYFP expression and location of the fibre optic cannulae for rats included in the analyses of visual threat. After histology, group sizes were: group Loom - eYFP n = 8, group Loom - Halo n = 8, group Control – Halo n = 3.

Figure 1F shows the mean and SEM velocities of rats in the 2 s pre-loom period, the 0.6 s looming period and the 10 s post loom, averaged across the three presentations of the looming stimulus. Animals showed an initial increase then decrease in velocity during these trials so that there was a significant quadratic trend in velocity (F (1, 15) = 4.77, p = .045). However, there was neither an overall difference between eYFP and Halo groups (F (1, 15) = 2.26, p > .05) nor an interaction between groups and time period (F (1, 15) < 1, p > .05). So, optogenetic inhibition of VLPAG had no effect on the locomotor response to a looming threat. Optogenetic inhibition did affect expression of post-encounter defensive behaviour in the two minutes following presentations of the looming stimulus (Figure 2F). The Halo group showed significantly more freezing compared to the eYFP group and to the control Halo group that received identical photoinhibition of VLPAG but in the absence of any looming stimulus (F (1, 15) = 7.54, p < .0001), and the two control groups did not differ from each other (F (1, 15) < 1, p > .05). Analysis of post-encounter defense across the three looming trials yielded a similar conclusion (Figure 2F). There was no freezing prior to the first presentation of a looming stimulus and there was significantly greater freezing in the Halo group compared to the two controls F (1, 15) = 7.54, p < .0001) which did not differ from each other (F (1, 15) < 1, p > .05). There was not a significant increase in freezing across trials, averaged across groups (F (1, 15) = 2.42, p > .05) and there were no interactions between group and trials F (1, 15) < 3.02, p > .05).

## Discussion

We examined the effects of brief optogenetic inhibition of LPAG and VLPAG neurons during US delivery on the acquisition of Pavlovian fear conditioning. Silencing LPAG or VLPAG neurons during the shock US augmented acquisition of contextual fear learning compared to a control eYFP group and a control group receiving photoinhibition offset from the US. A similar augmentation of learning was observed in auditory cue fear conditioning. Brief silencing of LPAG or VLPAG also increased expression of freezing as a post-encounter defensive behaviour to a visual threat (De Franceschi et al., 2016; Wei et al., 2015; Yilmaz & Meister, 2013) without affecting the locomotor response to this threat and without eliciting defensive behaviour itself in the absence of threat. These results support the claim that in addition to its role in expression of defensive behaviour (Assareh et al., 2016; Carrive & Morgan, 2003; Fanselow & Lester, 1988), L/VLPAG serve a critical role in fear learning (Fanselow, 1998; McNally et al., 2011; McNally & Westbrook, 2006), and show that this role is linked to the time of aversive US delivery.

There have been two coherent accounts of the role of PAG in fear learning. First, VLPAG controls a conditioned analgesic response that dampens or impedes detection of the shock US (Bolles & Fanselow, 1980; Fanselow, 1998). This conditioned analgesia reduces the impact of the shock US across conditioning trials and so reduces fear learning. Manipulations that reduce this analgesic response should permit continued detection of the shock US across conditioning and so augment fear learning. It is possible that optogenetic inhibition of LPAG/VLPAG during the shock US reduced analgesia and so augmented fear learning via this mechanism. However, this same optogenetic inhibition also augmented expression of freezing as a post-encounter defensive response to a non-noxious threatening visual stimulus. It is possible, even likely, that the visual threat was able to recruit pain modulatory mechanisms. However, there was no painful or noxious stimulus for such pain modulation to dampen and for optogenetic inhibition to augment. This is inconsistent with a pain modulatory mechanism being the critical mechanism through which PAG regulates fear learning. Indeed, other work also shows that PAG manipulations alter fear learning in the absence of a noxious US (Cole & McNally, 2008; McNally, Lee, Chiem, & Choi, 2005; McNally, Pigg, & Weidemann, 2004b; Parsons, Gafford, & Helmstetter, 2010). Moreover, PAG neurons that regulate fear learning are separate to those controlling pain modulatory responses (Ozawa et al., 2016).

Second, VLPAG regulates fear learning because it contributes to computation of a fear prediction error signal: the difference between the actual, *λ*, and expected outcomes, ΣV, of a conditioning trial (Cole & McNally, 2007; 2008; McNally et al., 2011; McNally, Pigg, & Weidemann, 2004a). Specifically, VLPAG neurons encode the expected outcome, ΣV, of a conditioning trial. **Σ**V increases across the course of conditioning to reduce the magnitude of prediction error, (*λ* - **Σ**V) (Rescorla & Wagner, 1972) and so constrain learning. It follows that optogenetic inhibition of these neurons at the time of reinforcement should artificially decrease ΣV, hence maintaining prediction error across conditioning and augmenting fear learning. Our results are consistent with this. Optogenetic inhibition of LPAG/VLPAG at the time of US delivery augmented contextual and auditory cue fear conditioning based on a noxious footshock US and also augmented development of freezing as a post-encounter response to a non-noxious looming visual stimulus.

Silencing either LPAG or VLPAG neurons during the US had the same facilitatory effect on learning, suggesting functional overlap between these distinct PAG columns in fear learning. This is reminiscent of past work using microinjections that also demonstrated overlap between these PAG columns in fear learning (Cole & McNally, 2007; 2008; McNally & Cole, 2006), but it contrasts with the pronounced differences in topography and properties of defensive behaviours controlled by the LPAG and VLAG (Assareh et al., 2016; Carrive, 1993; Fanselow, 1991; Vianna, Graeff, Brandao, & Landeira-Fernandez, 2001). The reasons for these differences in PAG columnar contributions to the acquisition versus expression of Pavlovian fear learning, and how they relate to the specific anatomical connectivity of LPAG and VLPAG remain to be determined. Regardless, this difference underscores the distinct contributions of PAG to the acquisition and expression of fear learning.

In summary, we show that LPAG and VLPAG serve a key role in regulation of Pavlovian fear learning at the time of US delivery. This role is parsimoniously explained by existing models which state that PAG contributes to computation of a fear prediction error signal that determines variations in the effectiveness of the aversive US in supporting learning.

## Acknowledgements

These experiments were supported by a grant from the National Health and Medical Research Council (1077806) and a Future Fellowship from the Australian Research Council (FT120100250).

